# Pharmacological Restoration of Mitochondrial Permeability Transition Pore Flickering by High-Content Drug Repurposing

**DOI:** 10.64898/2026.01.23.701330

**Authors:** Emanuela Franchini, Matteo Bulloni, Anna Sorgente, Katarina Paulikova, Ilaria Marafelli, Irene Sambri, Linda Pattini, Giorgio Casari

## Abstract

The mitochondrial permeability transition pore (mPTP) is a voltage- and calcium-regulated channel located in the inner mitochondrial membrane whose activity critically influences cellular fate. While prolonged pore opening leads to mitochondrial depolarization, matrix swelling, and cell death, brief and reversible opening events, referred to as flickering, enable controlled release of calcium and reactive oxygen species and serve essential physiological functions. Emerging evidence indicates that restoring physiological mPTP flickering, rather than suppressing pore activity, may be beneficial in disorders characterized by impaired pore dynamics, including hereditary spastic paraplegia type 7 (SPG7). However, no approved therapies are currently available to promote controlled mPTP pore opening.

To identify pharmacological modulators of flickering, we performed a high-content screening of 2,000 FDA and EMA-approved compounds using a validated fluorescence-based assay coupled with automated image analysis. Thirteen compounds increased both the frequency and the area of flickering events while preserving cellular and mitochondrial integrity. Validation in fibroblasts derived from two SPG7 patients and healthy controls confirmed reproducible activity across distinct genetic backgrounds. Among the prioritized candidates, berberine emerged as the most robust modulator, consistently enhancing mPTP flickering independently of SPG7 mutation status. Notably, berberine selectively increased the proportion of small-size flickering events, indicative of physiological pore activity. These findings identify berberine as a promising modulator of mPTP dynamics and support pharmacological restoration of physiological flickering as a potential therapeutic strategy for SPG7 and other disorders associated with impaired mitochondrial permeability transition pore regulation.

## Introduction

The mitochondrial permeability transition pore (mPTP) is a voltage- and calcium-sensitive channel in the inner mitochondrial membrane [1]. Sustained mPTP opening leads to mitochondrial membrane potential dissipation, matrix swelling, and cell death, contributing to pathological conditions including ischemia-reperfusion injury and neurodegenerative diseases [2]. In contrast, transient mPTP opening events, termed “flickering”, allow controlled calcium and reactive oxygen species (ROS) efflux without triggering cell death, functioning as a mitochondrial “safety valve” [3] and establishing it as an essential physiological process distinct from pathological pore opening [4].

Emerging evidence demonstrates that flickering dynamics are altered during aging, with senescent cells exhibiting increased flickering as an adaptive mechanism to manage elevated mitochondrial calcium, [5]. This finding supports the notion that mPTP flickering represents an adaptive mitochondrial process whose loss during aging may become maladaptive, contributing to impaired calcium and redox homeostasis.

Spastic paraplegia type 7 (SPG7), caused by mutations in the *SPG7* gene encoding the mitochondrial protease paraplegin [6], provided the first evidence that impaired flickering could drive human disease, as shown in SPG7 patient fibroblasts and in neuronal cells of the *Spg7 K.O* mouse model [7]. Treatment with Bz-423, a benzodiazepine favoring the mPTP opening and available only for research purpose, restored flickering in SPG7 cells and improved motor function in *Spg7^-/-^* mice [7]. Unfortunately, Bz-423 use is limited to experimental applications and has the potential to trigger apoptosis [8].

Drug repurposing offers a rapid path to clinical translation by leveraging established safety and pharmacokinetic data [9]. FDA and EMA-approved mPTP agonists, if present, could be used off-label in SPG7 patients while enabling mechanistic studies of flickering modulation. Beyond SPG7, such compounds could address flickering deficits in neurodegenerative diseases where mitochondrial dysfunctions are central [10].

Here, we screen a library of 2,000 FDA/EMA-approved compounds using a quantitative double fluorescence-based flickering assay in SPG7 patient fibroblasts. We identify 13 candidates and validate them across multiple genetic backgrounds. We finally prioritize berberine, a natural alkaloid [11], as a lead mPTP agonist with translational potential for SPG7 and other mitochondriopathies.

## Material & Methods

### Cell cultures

Human primary control fibroblasts and *SPG7*-*A* fibroblasts were cultured in Dulbecco’s Modified Eagle Medium (DMEM; Cat. #ECM0728L, EuroClone, Milan, Italy) containing 10% fetal bovine serum (FBS; Cat. #35-079-CV, Corning®, Corning Incorporated, NY, USA) and 1% antibiotics (penicillin/streptomycin; PS; Cat. #ECB3001D, EuroClone, Milan, Italy) at 37 °C and 5% CO_2_. All cells were split upon reaching confluence, by rinsing the monolayer twice with sterile Dulbecco’s Phosphate-Buffered Saline (DPBS; Cat. #21-031-CV, Corning®, Corning Incorporated, NY, USA) without calcium and magnesium and exposing cells to 0.25% Trypsin-EDTA (Cat. #ECM0920D, EuroClone, Milan, Italy) for 2-3 min at 37°C. Detached cells were centrifuged to remove Trypsin from the media and then replated in fresh new medium.

### Visualization and quantification of TMRM flickering events

Our approach allows the measurement of transient mitochondrial membrane potential changes resulting from mPTP opening, visualized as flickers of fluorescence. To assess flickering events, we used two different fluorescent dyes: Tetramethylrhodamine (TMRM Cat #I34361, Thermo Fisher Scientific, Waltham, MS, USA) and MitoTracker Green (MTG Cat #M7514, Thermo Fisher Scientific, Waltham, MS, USA). TMRM is a rapidly repartitioning cationic, red-orange, fluorescent dye that is readily sequestered by polarized mitochondria, whereas MTG is a cell-permeant, green-fluorescent mitochondrial dye whose entrance is not dependent on mitochondrial membrane potential (ΔΨm) and is used as a good marker for mitochondrial morphology. Cells were plated on µ-Slide 8 well high ibiTreat (Cat #80806, Ibidi GmbH, Gräfelfing, Germany), and treated with a complete Dulbecco’s Modified Eagle Medium without phenol red (Cat #31053044, Thermo Fisher Scientific, Waltham, MS, USA) containing 10% FBS), 20 nM TMRM, 2µM cyclosporine H (CsH) (Cat# SML1575, Sigma-Aldrich, St. Louis, MS, USA) and 100 nM MTG for 30 min at 37°C. Under these conditions TMRM operates in non-quenching mode. Cells were then washed twice with DPBS, and the slide was placed on the stage of GE Healthcare DeltaVision microscope (GE Healthcare, Waukesha, WI, USA) equipped with a stage-top incubator and CO_2_ control system. Loss of TMRM induced by mPTP opening was quantified from time-lapse image stacks (1s intervals, 200 ms exposure) acquired over approximately 5 min using with a 60X oil-immersion objective. Laser power was attenuated to 2% to minimize photobleaching and phototoxicity. Imaging was performed at the Advanced Light and Electron Microscopy BioImaging Center (ALEMBIC), IRCCS San Raffaele Hospital, Milan, Italy.

### mPTP activity detection by the Matlab pipeline

Flickering events were detected sequentially using a custom MATLAB script that processes the two-channel frame sequences of each video. The green channel, reporting MTG dye intensity, was used to delineate the boundaries of the mitochondrial network and calculate its total area. The red channel, reflecting TMRM intensity, was analyzed to identify flickering events by computing the frame-to-frame fluorescence difference (TMRM(n+1) − TMRM(n)).

The automated pipeline quantified the mitochondrial area involved for each event (*relative flickering area (RFA*) and the overall frequency of events throughout the time-lapse (*flickering frequency (FF)*.

### HCS setting conditions

*SPG7*-*A* fibroblasts were seeded at 15,000 cells per well through automatization using robotic liquid handle (JANUS® G3 Automated Workstation, PerkinElmer, Waltham, MA, USA) at the High-Throughput Screening (HiTS) Facility, Department of Biology, University of Padova (Padova, Italy) in a 96-wells plate (96 Well Polystyrene µClear®, Cat #655090, Greiner Bio-One, Kremsmünster, Austria) to minimize operator variability. The automated protocol encompassed cell seeding, drug library preparation, compound and dye treatments, washing steps, and medium addition. Following overnight incubation at 37 °C with 5% CO_2,_ cells were pharmacologically treated for 2h (at 37°C with 5% CO_2_) with drugs of the library (L1021-DiscoveryProbe-FDA-approved-Drug-Library, APExBIO, Houston, TX, USA) dissolved in dimethyl sulfoxide (DMSO) and positive (Bz-423, Cat# SML1944-5 mg, Sigma-Aldrich, St. Louis, MS, USA) and negative (DMSO, Cat#D4540-100ML, Sigma-Aldrich, St. Louis, MS, USA) controls at a final concentration of 500 nM. To prevent cell detachment during solution exchanges, all liquid handling was performed at 10 µL/s dispensing velocity with pipette tips positioned 3 mm above the well bottom, as optimized in preliminary experiments. Drug removal was performed through three sequential aspiration cycles (50 µL each), followed by a washing step with 150 µL DPBS. After DPBS removal, cells were incubated for 30 min at 37 °C, protected from light, with complete DMEM w/o phenol red, containing 20 nM TMRM, 2 µM CsH, 100 nM MTG, and the individual drugs from the library. After staining solution removal, cells were washed once more in DPBS before adding fresh DMEM without phenol red containing CsH and the respective drugs. Images were acquired with Operetta High Content Screening System (PerkinElmer, Waltham, MA, USA) using a 40X water-immersion objective. Two fields per well were acquired with 110 time points per field. For each field, images were acquired at 2-second intervals over a total acquisition time of 3 min 40 s, with illumination times of 50 ms for the green channel and 20 ms for the red channel, using 100% light power. Data analysis was performed using a MATLAB Script (MathWorks, Natick, MA, USA).

### HCS-Based Candidate Selection Criteria

Compounds were classified as positive hits based on a normalized flickering rate threshold of 0.01 events s^-1^μm^-2^, calculated as the ratio between the number per second of absolute flickering events and total mitochondrial area. This empirical cut-off was set to balance sensitivity and specificity in candidate selection. Lower thresholds would have yielded excessive false positives, making subsequent validation impractical, while higher thresholds risked excluding compounds with moderate but potentially therapeutically relevant activity.

All “flick-positive” compounds were cataloged in a comprehensive database documenting: plate position, CAS number, therapeutic category, targeted biological pathway, known molecular target, clinical indication, and normalized flickering metrics (FF and RFA). Compound annotations were retrieved from DrugBank [12] and the IUPHAR/BPS Guide to Pharmacology [13].

### Cellular and mitochondrial morphological analysis for drug selection

The morphological analysis of cells after treatment was performed using Volocity 3D Image Analysis Software (Quorum Technologies Inc., Puslinch, ON). This method was designed to visualize the impact of each compound on cellular morphology and to exclude those causing pronounced alterations indicative of cytotoxicity. Volocity® is an image analysis platform capable of processing both 2D and 3D microscopy datasets. In this study, an automated analysis pipeline was employed to identify and quantify the number of mitochondria and cells within each image. The software performs mitochondrial compartmentalization at the single-cell level and computes several morphological parameters, including cell perimeter, shape factor, and maximum cell length (indicator of cell elongation). All parameters are exported to table (Supplementary Fig. 1). The shape factor, a dimensionless parameter ranging from 0 to 1, was calculated to quantify how closely the cell shape resembles a perfect circle (in 2D) or sphere (in 3D). A value of 1 indicates perfectly round or spherical shape, whereas lower values correspond to more elongated forms. For mitochondrial network analysis, Mitochondria Analyzer [14] was employed to generate automated masks from fluorescence images and calculate morphological parameters at both cellular and subcellular levels. Three representative timepoints per video (beginning, midpoint, and end of incubation) were analyzed to capture temporal dynamics. Critical assessment parameters included mitochondrial number per cell, total areas, branch connectivity metrics, and mean form factor, a key indicator for fragmented, rounded mitochondria, or elongated morphology (Supplementary Fig. 1). Form factor values approaching 1.0 indicate fragmented, rounded mitochondria, while higher values reflect elongated morphology.

Furthermore, a toxicity score ranging from 0 to 2 was assigned to each candidate based on morphological assessment. A score of 0 indicated no detectable alterations in mitochondrial and cellular morphology, while 2 denoted compounds affecting both cellular morphology and mitochondrial network integrity. Scores of 1 identified drugs impacting only one parameter, and 0.5 was assigned to candidates exhibiting mild effects on either mitochondrial fragmentation or cellular shape.

### Pharmacological treatment with the 13 selected compounds

Wild-type human fibroblasts and *SPG7-A* patient fibroblasts were seeded at 15,000 cells per well in black-walled 96-well plates (to avoid photobleaching of the nearby wells) in 150 µL of DMEM supplemented with 10% FBS. Cells were treated with the 13 selected compounds: adapalene (Cat#A7486-10MG, Sigma-Aldrich, St. Louis, MS, USA); eriodictyol (Cat#94258, Sigma-Aldrich, St. Louis, MS, USA)-5MG; (R) -(+)- lipoic acid (Cat#07039-10MG, Sigma-Aldrich, St. Louis, MS, USA); lumiracoxib (Cat#SML2928-10MG, Sigma-Aldrich, St. Louis, MS, USA), ketoprofen (Cat#K1751-1G, Sigma-Aldrich, St. Louis, MS, USA), vildagliptin (Cat#SML2302-50MG, Sigma-Aldrich, St. Louis, MS, USA); ezetimibe (Cat# SML1629-25MG, Sigma-Aldrich, St. Louis, MS, USA); CP-945598 (Cat#PZ0019-5MG, Sigma-Aldrich, St. Louis, MS, USA); biotin (Cat#B4501-100MG, Sigma-Aldrich, St. Louis, MS, USA); 2-deoxy-D-glucose (Cat# D8375-10MG, Sigma-Aldrich, St. Louis, MS, USA); 4-aminopyridine (Cat#A78403-25G); uridine (Cat#U3003-5G, Sigma-Aldrich, St. Louis, MS, USA); and berberine hydrocloride (Cat#004CA10006427-10, Cayman Chemical, Michigan, USA) at 500 nM concentration or respective vehicle controls for 2 hours at 37°C with 5% CO₂. adapalene, eriodictyol, lipoic acid, lumiracoxib, ketoprofen, vildagliptin, ezetimibe, CP-945598, and berberine were dissolved in DMSO, according to their solubility characteristics. biotin, 2-deoxy-D-glucose, 4-aminopyridine, and uridine were dissolved in water. Cells were washed twice with DPBS and incubated with the staining solution containing complete DMEM without phenol red supplemented with 10% FBS, 20 nM TMRM, 2 µM CsH, 100 nM MTG, and 500 nM test compounds for 30 minutes at 37°C, protected from light. The staining solution was removed, cells were washed twice with DPBS, and fresh DMEM without phenol red containing 10% FBS, 2 µM CsH, and 500 nM compound was added. Images were acquired using the ImageXpress HCS High-Content Screening System (Molecular Devices, LLC, San Jose, CA, USA) and EVOS microscope equipped with an incubator and CO₂ control system using 40X water immersion objective. 240 time points were acquired for each field at 1-second intervals, with 80 ms illumination time for green and red channels (4-minute acquisition period per field). Each well was imaged at four distinct sites to prevent photobleaching. Flickering events were quantified using a custom MATLAB script (MathWorks, Natick, MA, USA).

### Statistical analysis

Statistical comparisons between two groups were performed using unpaired Student’s t-test or Mann-Whitney U test when normal distribution could not be assumed. Flickering data generated by the MATLAB script from each experiment were normalized to the mean of control cell samples or vehicle-treated controls and are reported as scatter dot plots with median lines. Statistical analyses were performed using GraphPad Prism 10. Data were tested for normality and homogeneity of variance using the Shapiro-Wilk normality test. A p-value <0.05 was considered statistically significant and indicated by an asterisk. All experiments were performed independently at least three times under identical conditions.

## Results

### Characterization of mitochondrial flickering events

Leveraging our previous findings demonstrating impaired transient mPTP opening in SPG7 condition [7] we refined and strengthened the biological assay to ensure reliable detection and quantification of mitochondrial flickering events.

To this end, we employed two complementary fluorescent probes: TMRM, a cationic red–orange dye that accumulates in polarized mitochondria, and MTG, a green fluorescent mitochondrial marker which fluorescence is independent of ΔΨm [15].

Under physiological mitochondrial membrane potential, both fluorescent dyes are taken up by mitochondria. The overlap of red TMRM and green MTG signals produces a yellow-labelled mitochondrial network. Upon mPTP opening, TMRM rapidly redistributes from the release site, resulting in a transient local increase followed by a sharp decrease in red fluorescence intensity, which shifts the mitochondrial staining to green. The minimum red fluorescence intensity corresponds to the maximal release of TMRM during the flickering event. Once the mPTP closes, the ΔΨm is restored, allowing TMRM to re-enter mitochondria and leading to a rise in TMRM fluorescence. After closure, the initial fluorescence levels are reestablished, and the mitochondrial network returns to its original yellow appearance (Fig. 1).

**Figure 1.**
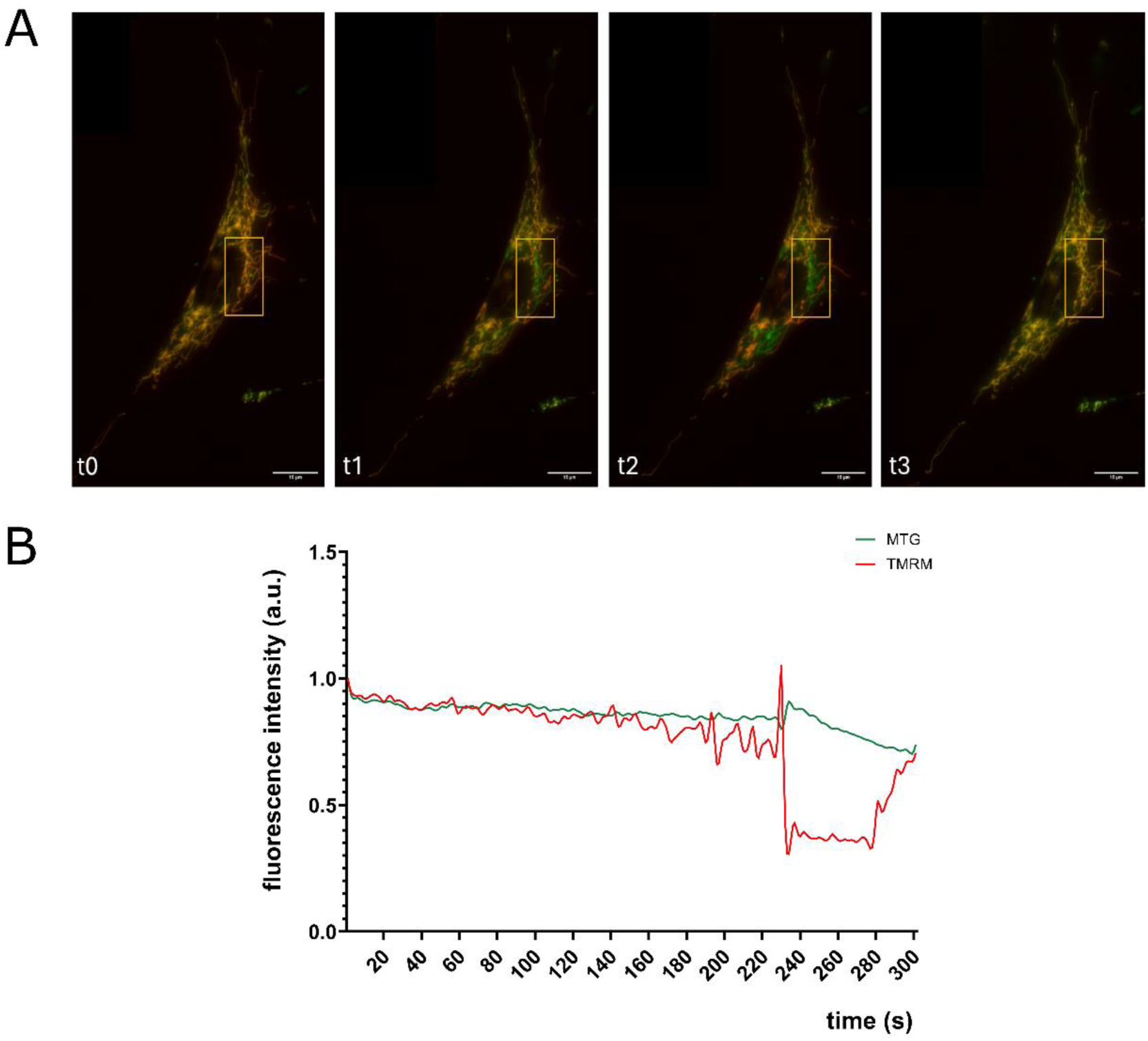
Mitochondrial flickering visualization using TMRM and MTG dyes in control fibroblasts. **(A)** Sequential imaging frames captured during video microscopy with GE Healthcare DeltaVision system showing wild-type primary fibroblasts and documenting the temporal progression of a flickering event. Scale bar =15 µm **(B)** Graphical representation of a single flickering event recorded over 300 seconds (s). MTG: MitoTracker Green, TMRM: Tetramethylrodamine.

### Automated Video processing for flickering detection, quantification and analysis

To enable automated, quantitative detection of flickering events through combined spatial and temporal analyses, we developed a MATLAB-based custom script. Within this pipeline, the green channel is used to segment the mitochondrial network and calculate its total area (Fig. 2 A); subsequently, each video frame is analyzed to detect individual flickering events. An mPTP flickering event is detected only when all criteria listed below are simultaneously satisfied within a minimum-size region of interest (hereafter referred to as the “event area”) of 4 μm^2^, ensuring high specificity and minimizing false positives due to mitochondrial movement. These criteria are: (i) the mean TMRM intensity in the event area must be above a minimum threshold for three seconds, to avoid false positives in background regions; (ii) the average change in TMRM intensity between the current and previous frame in the event area must be negative and lower than a minimum threshold, indicating a rapid TMRM drop; (iii) the decrease in TMRM intensity must exceed the corresponding MTG decrease signal multiplied by a predefined shrinking factor, excluding global fluorescence fluctuations.; (iv) no more than 35% of the event area may exhibit a *positive* increase in TMRM intensity, to avoid detecting shifting artifacts rather than true events and (v) TMRM intensity must recover within the 15 seconds following a putatively identified flickering event, consistent with transient pore opening followed by membrane repolarization. Detected events occurring in close spatial or temporal proximity are subsequently merged to avoid duplicate counting of the same biological event. Ultimately, for each video, the automated pipeline quantifies two parameters:

1. *flickering frequency (FF)*: the overall frequency of events per mitochondrial network area, computed as: (number of flickering events / time-lapse duration) / total mitochondrial area;
2. *relative flickering area (RFA)*: the percentage of mitochondrial area undergoing flickering events per second, computed as: (total flickering area throughout time-lapse / time-lapse duration) / total mitochondrial area).

**Figure 2.**
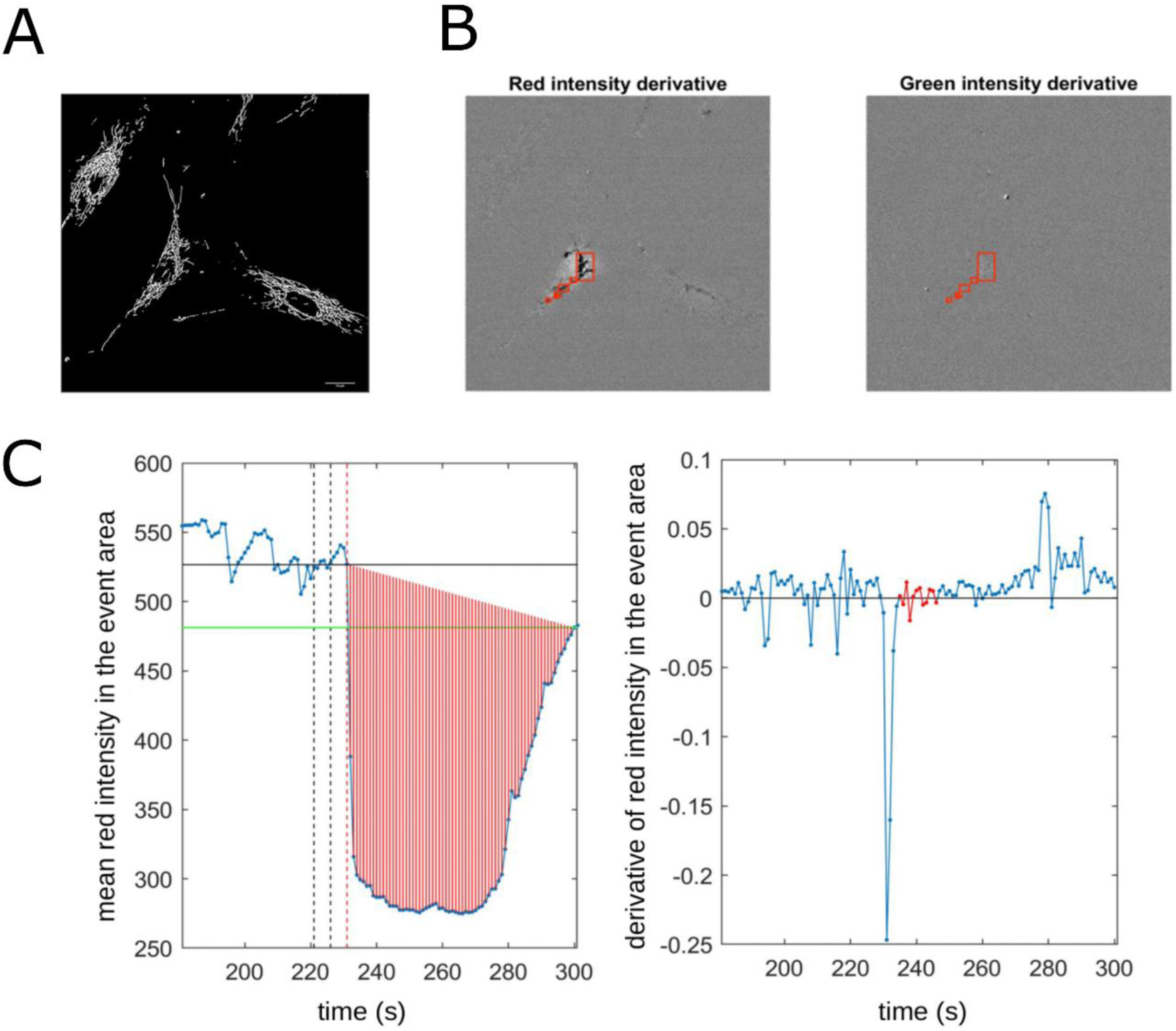
Example output of the video processing pipeline for flickering events identification. **(A)** Binary mask of the mitochondrial network generated from the green (MTG) channel. Scale bar =15 µm. **(B-C)** Example graphical output yielded by the pipeline for each individual event identified. **(B)** Spatial identification of the event via intensity derivatives in the red (TMRM) and green channels. **(C)** Temporal profile of the detected flicker: the left graph plots the mean TMRM intensity in the event area, illustrating the fluorescence drop and recovery (red shaded region) in time, while the right graph displays the derivative of the mean TMRM intensity in the event area, highlighting the negative spike associated with the rapid TMRM release at event occurrence.

Figures 2B and 2C show the graphical outputs generated using a custom MATLAB script.

### High Content Screening for identification of mPTP flickering modulators

To identify the most suitable cellular substrate for screening, we chose newly established *SPG7-A* fibroblasts, which have a low passage number, underwent minimal *in vitro* manipulation, and carry compound heterozygous mutations (p.Gly349Ser and p.Leu706Glnfs). As expected, *SPG7-A* exhibited a marked reduction in flickering parameters compared with wild-type fibroblasts (Fig. 3, B).

**Figure 3.**
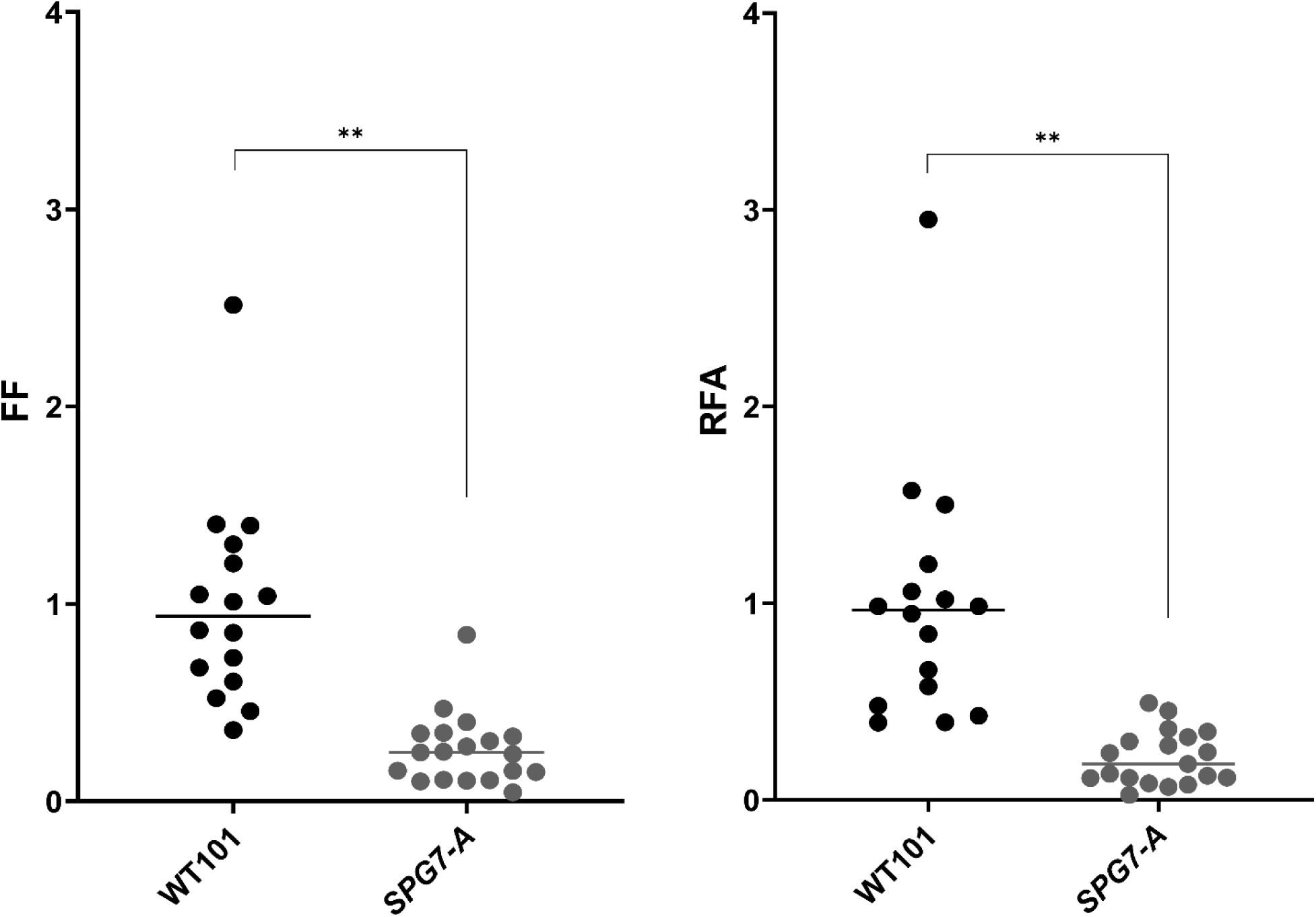
Flickering impairment in SPG7 patient fibroblasts vs WT human fibroblasts. Scatter dot plots of 2 parameters describing FF and RFA for WT (black) and *SPG7-A* (grey). Four independent experiments; Mann Whitney p value ** < 0.0001.

A High Content Screening (HCS) campaign was performed using a library of approximately 2,000 chemical compounds with the goal of identifying novel mPTP agonists with favorable safety profiles. All compounds, together with Bz-423 as positive control, were applied at a final concentration of 500 nM to maximize sensitivity while minimizing the risk of false negatives due to under-dosing.

Following HCS acquisition, the resulting dataset was processed using the MATLAB-based analysis pipeline described above. The results were then summarized in a heatmap that displays the effect of each compound, normalized to the total mitochondrial network area. This visualization enables rapid and straightforward identification of candidate molecules capable of inducing mPTP opening (Fig. 4A and 4B). A white-to-blue color gradient highlights compounds that increase flickering activity in terms of both FF and RFA, with darker blue indicating positive candidates and white indicating no detectable effect.

**Figure 4.**
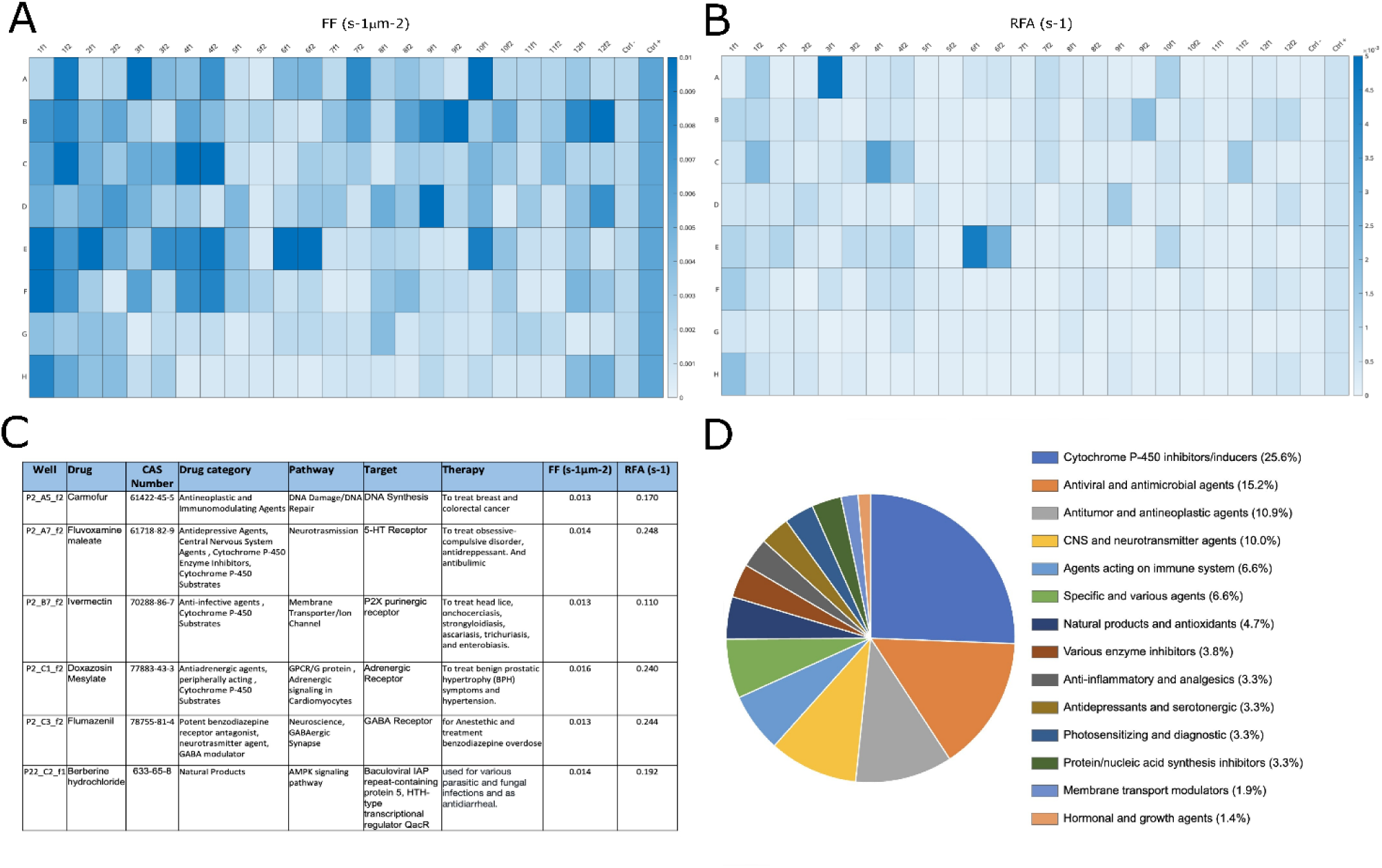
High-throughput screening results displayed as 96-well plate heatmaps illustrating two quantitative flickering parameters. **(A)** FF and **(B)** RFA. Each well contains a distinct compound, with positive and negative controls located in the rightmost column. Two representative imaging fields were analyzed per well. In both panels, the color scale allows rapid visualization of compounds with increased flickering activity, with higher intensity indicating greater FF or RFA. **(C)** Representative examples of compounds identified as promising flickering enhancers based on combined FF and RFA parameters. **(D)** Classification of the 91 flickering-positive compounds according to their therapeutic categories.

Based on the predefined selection criteria, 91 compounds that increased flickering activity were initially identified as positive hits with representative examples shown in Fig. 4C and a graphical overview of their drug categories presented in Fig. 4D. Although no single pharmacological class predominated, the most frequently represented categories included cytochrome P450 modulators, antimicrobial agents, antineoplastic drugs, central nervous system–targeting compounds, and immunomodulatory agents.

### Rational prioritization of candidate mPTP agonists

We applied a rational prioritization strategy based on three complementary approaches to refine the list of positive hits, excluding compounds associated with cellular toxicity or pharmacological properties unsuitable for long-term therapeutic use.

The first prioritization step involved quantitative analysis of cellular morphology, including shape factor and maximum length. Under toxic conditions, cells typically lose their elongated morphology and adopt a more rounded shape, a clear hallmark of early cell death or distress. An increase in shape factor, together with a reduction of the longest cell axis, is therefore suggestive of a toxic insult.

Following cell morphology analysis, mitochondrial network integrity was evaluated to assess organellar toxicity. Critical assessment parameters included mitochondrial number per cell, total areas, branch connectivity metrics, and mean form factor, a key indicator for fragmented, rounded mitochondria or elongated mitochondrial morphology (Supplementary Fig. 1).

A third level of evaluation focused on pharmacological properties relevant to chronic administration, prioritizing compounds with good patient accessibility, well-established long-term safety profiles, and simple routes of administration. Antineoplastic agents were excluded because their non-selective mechanisms and severe adverse effects make them unsuitable for prolonged treatment.

The compounds that passed this multiparametric selection were mainly antioxidants, potentially beneficial for mitigating the excessive oxidative stress characteristic of the SPG7 condition, natural products with favorable safety for long-term use, and agents offering dual benefits by reducing mitochondrial oxidative stress while promoting mitochondrial biogenesis. Blood-brain barrier permeability was also considered essential, given that the therapeutic target lies within the central nervous system.

This multiparametric classification strategy reduced the initial pool of hits to 13 high-priority candidates, subsequently grouped into three functional categories based on their primary mechanisms of action (Table 1).

**Table 1.** Prioritized therapeutic candidates. Thirteen compounds were identified through systematic classification analysis. For each candidate, the table reports: therapeutic class, current clinical indication, pathway involvement, molecular target, route of administration, toxicity score (0–2), in vitro concentration range, and an integrated benefit–risk profile.

| Drug | Drug Category | Therapeutic Application | Pathway | Target | Route of Administration | Toxic Effect (0-2) | Benefit | Risk |
| --- | --- | --- | --- | --- | --- | --- | --- | --- |
| Berberine hydrochloride | Alkaloids, natural products | used for various parasitic and fungal infections and as antidiarrheal | AMPK signaling pathway | Baculoviral IAP repeat-containing protein 5, HTH-type transcriptional regulator QacR | Oral | 0 | Orally administered compound with well-documented safety profile. Clinically employed for treating diabetes, hyperlipidemia, and cardiovascular diseases. Exhibits antimicrobial and neuroprotective properties, demonstrating efficacy in murine models of multiple CNS pathologies (Alzheimer's disease, Parkinson's disease, cerebral ischemia, depression, schizophrenia, epilepsy) through oxidative stress reduction, anti-inflammatory activity, and cerebral damage prevention. | High doses inhibit cell proliferation and induce apoptosis |
| Uridine | Nucleic acid precursor, Nucleotide/nucleoside supplement | Management of neuropsychiatric deficits associated with cerebrovascular disease in combination with cytidine | DNA synthesis pathway | U6 snRNA-associated Sm-like protein LSM6, Nucleoside-specific channel-forming protein tsx | Oral | 0 | Oral bioavailability. Promotes mitochondrial biogenesis and mitophagy in chronic diabetic mouse models, attenuates apoptosis and oxidative stress, reduces Iba1+ microglia activation in Alzheimer's disease models, and activates PGC-1α. | No significant impact on calcium retention capacity or potassium transport mechanisms |
| Ezetimibe | Cholesterol absorption inhibitor, Cytochrome P-450 enzyme inhibitor | Reduction of total cholesterol, LDL-cholesterol, apolipoprotein B, and non-HDL cholesterol in primary hyperlipidemia and familial hypercholesterolemia | Cholesterol absorption pathway | Niemann-Pick C1-like protein 1, Sterol O-acyltransferase 1, Aminopeptidase N | Oral and intravenous | 1 | Oral administration; activates AMPK, SIRT1, and PGC-1α in Parkinson's disease rat models; upregulates Nrf2 thereby reducing oxidative stress; synergistic effects with simvastatin in hyperlipidemia treatment; blood-brain barrier penetration; excellent safety profile | — |
| DL-α-lipoic acid | Decarboxylation cofactor, Antioxidant compound | — | Endocrine and hormonal pathways | — | Oral | 0 | Oral bioavailability; ameliorates mitochondrial oxidative stress; frequently combined with L-carnitine; demonstrated efficacy across multiple animal models; pharmacokinetic profile characterized in canine studies | Dose-dependent adverse effects at elevated concentrations |
| Vildagliptin | Antihyperglycemic agent, DPP-4 inhibitor | Type 2 diabetes mellitus management | Glucose homeostasis regulation and glycemic control | Dipeptidyl peptidase-4 (DPP-4) | Oral | 1 | Oral administration; reduces reactive oxygen species production and ameliorates mitochondrial dysfunction in diabetic mouse models | — |
| Eriodictyol | Natural flavonoid compound | Neuroprotection, cardioprotective activity, hepatoprotective effects, anti-diabetic and anti-obesity properties, dermatoprotective effects with pronounced analgesic properties, antioxidant and anti-inflammatory activities, antipyretic and antinociceptive actions, antineoplastic activity | PI3K/mTOR signaling pathway, Flavonoid biosynthesis pathway, PI3K/Akt pathway | Peroxisome proliferator-activated receptor γ2 (PPARγ2) | Experimental/Nutritional supplementation | 2 | Inhibits ferroptosis in ischemic stroke through free radical scavenging; modulates SIRT1 in Alzheimer's disease; demonstrated neuroprotective, cardioprotective, hepatoprotective, anti-diabetic, anti-obesity, dermatoprotective, analgesic, antioxidant, anti-inflammatory, antipyretic, and antinociceptive properties | Investigational compound with limited clinical data |
| 4-aminopyridine | Membrane transport modulator, Cytochrome P-450 substrate, Potassium channel antagonist | Motor function enhancement in multiple sclerosis (MS) patients | Neuromuscular and synaptic transmission | Voltage-gated potassium channels | Oral | 0 | Oral administration with therapeutic dosing at 20-30 mg in divided doses over extended periods; significant improvement in ambulation, spasticity, and lower extremity function in MS patients; blood-brain barrier penetration | High mammalian toxicity profile causing hyperexcitability, salivation, tremors, muscular incoordination, clonic and tonic seizures, potential cardiac or respiratory arrest |
| CP-945598 HCl | Cannabinoid receptor antagonist | — | G-protein coupled receptor signaling | Cannabinoid receptor type 1 (CB1) | Under investigation | 0.5 | Enhances locomotor activity in murine models at 17.8 mg/kg dosing | Limited pharmacological data available |
| Biotin | Enzymatic cofactor, Vitamin B7 | Treatment of nail disorders | Gluconeogenesis, lipogenesis, fatty acid biosynthesis, propionate metabolism | Biotinylation enzymes, Propionyl-CoA carboxylase β-chain, Biotin-protein ligase, Methylcrotonyl-CoA carboxylase β-chain, Acetyl-CoA carboxylase 2 α-chain, Methylcrotonyl-CoA carboxylase α-subunit, Pyruvate carboxylase, Propionyl-CoA carboxylase α-chain, Acetyl-CoA carboxylase 1 | Oral | 0 | Enhances ATP production in oligodendrocytes *in vitro* | — |
| Lumiracoxib | Analgesic agent, Selective COX-2 inhibitor, Cytochrome P-450 substrate | Acute and chronic management of osteoarthritis symptoms in adults, particularly knee osteoarthritis | Prostaglandin synthesis inhibition | Prostaglandin G/H synthase 2 (COX-2) | Oral | 1 | Favorable tolerability profile with minimal adverse effects, lacking gastrointestinal and cardiac toxicity. | COX-2 inhibitors suppress ATP synthesis |
| Ketoprofen | Nonsteroidal anti-inflammatory drug (NSAID) with analgesic and antipyretic properties | Treatment of rheumatoid arthritis, osteoarthritis, ankylosing spondylitis, dysmenorrhea, mild-to-moderate musculoskeletal pain, postoperative pain, and postpartum pain | Prostaglandin synthesis inhibition | Prostaglandin G/H synthase 1 and 2 (COX-1/COX-2) | Oral, topical, intramuscular | 1 | — | Gastrointestinal adverse effects |
| Adapalene | Topical anti-acne agent, Anti-inflammatory compound | Treatment of acne vulgaris in adolescents and adults | Retinoic acid signaling pathway | Retinoic acid receptor RXR-β, Retinoic acid receptor RXR-γ, Retinoic acid receptor RXR-α, Transcription factor AP-1, Toll-like receptor 2, Aspartate aminotransferase | Topical (gel or cream formulation with transdermal absorption) | 0.5 | Topical route of administration with minimal adverse effects (cutaneous irritation). | Exhibits antiproliferative activity and promotes apoptotic cell death. |
| 2-deoxy-D-glucose | Anti-infective agent, Antimetabolite, Glucose analog | Metabolic assessment in cardiovascular disease, oncology, and Alzheimer's disease; antineoplastic and antiviral applications | G-protein coupled receptor signaling | Hexokinase | Oral | 0 | Oral administration; inhibits neoplastic cell proliferation and N-linked glycosylation; functions as glycolytic inhibitor and caloric restriction mimetic; inhibits pathogen-associated molecular patterns (PAMPs); anticonvulsant properties; activates PI3K/Akt/mTOR, Ras/Raf/MAPK, and PKC signaling cascades; stabilizes hypoxia-inducible factor-α (HIF-α); activates sirtuin-4 (SIRT4) | Inhibits glycolysis, tricarboxylic acid cycle, and pentose phosphate pathway resulting in ATP depletion, underlying its antineoplastic mechanism |

The first category comprises agents targeting oxidative stress and mitochondrial biogenesis, including: (i) berberine, an orally administered compound with well-documented safety profile, established clinical use for diabetes and hyperlipidemia, and cardiovascular diseases, and reported neuroprotective efficacy [16]; (ii) uridine, which was demonstrated to enhance mitochondrial biogenesis and mitophagy in diabetic mouse models [17]; (iii) ezetimibe, which upregulates Nrf2 expression and activates AMPK/SIRT1/PGC-1α signaling [18]; (iv) DL-α-lipoic acid; (v) vildagliptin; and (vi) eriodictyol, which similarly exhibited ROS reduction and neuroprotective effects in disease models [19,20].

The second category includes compounds that target motor dysfunction by enhancing neuromuscular transmission: (i) 4-aminopyridine, which improves ambulation and reduced spasticity [21]; (ii) CP-945598, a motor activity enhancer [22], and (iii) biotin, which stimulates ATP production in oligodendrocytes, potentially enhancing mPTP activity [23].

The third category comprises compounds with mixed activities that require careful benefit-risk evaluation, particularly in the context of chronic administration, including: (i) lumiracoxib, showing good tolerability but also causing mitochondrial ATP synthesis inhibition [24]; (ii) ketoprofen, with gastrointestinal adverse effects in chronic administration; (iii) adapalene, which showed antiproliferative activity requiring chronic safety validation [25], and (iv) 2-deoxy-D-glucose, which induces ATP depletion through glycolytic inhibition [26].

### Reconfirming drug candidates as flickering enhancers

The 13 prioritized compounds were subsequently tested in three human fibroblast cell lines to assess efficacy and consistency across different genetic backgrounds: the wild-type control (WT101), and the *SPG7-A* mutant fibroblasts.

As shown in Fig. 5, berberine demonstrated the most consistent and robust activity, significantly increasing both FF and RFA across all cell types. Ketoprofen significantly increased flickering area without affecting event frequency, suggesting a more limited or context-dependent effect.

**Figure 5.**
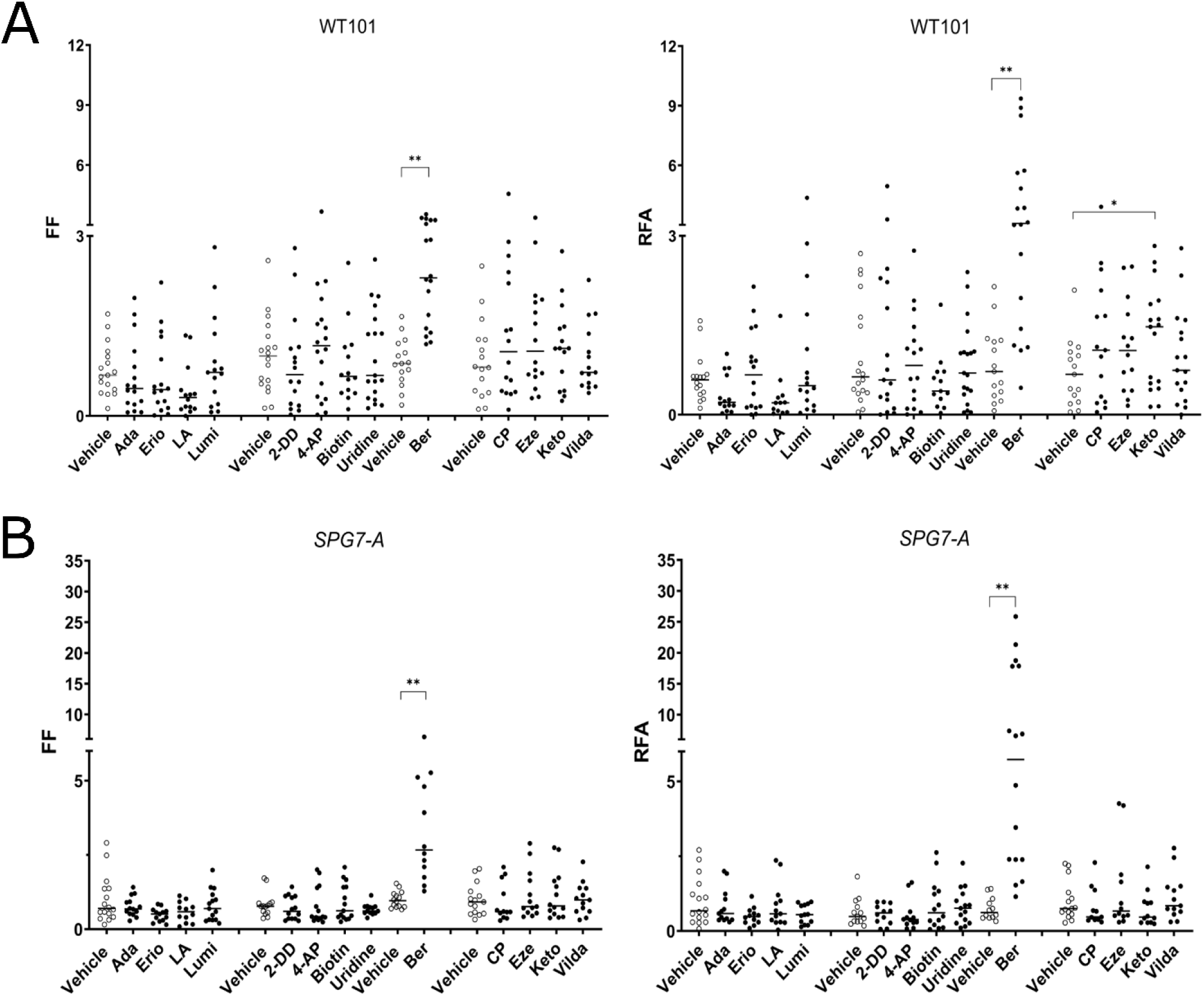
Biological assay validation of drug compounds performed in SPG7 patient fibroblasts and in WT human fibroblasts. Scatter dot plots of 2 parameters describing flickering events (FF and RFA) for **(A)** WT101 and **(B)** *SPG7-A* treated with selected drugs (black) and respective vehicle control (open circles). Four independent experiments for each sample are performed: (A)Mann Whitney ** p value <0.001 (FF); Mann-Withney ** p value< 0.0001 and Unpaired t test * p value < 0,05 (RFA). **(B)**; unpaired t test **p value <0.0001 (FF); Mann Whitney ** p value <0.0001 (RFA). Adaptalene (Ada); eriodictyol (Erio), lumiracoxib (Lumi); DL-α-lipoic acid (LA); 2-deoxy-D-glucose (2-DD); 4-aminopyridine (4-AP); berberine hydrocloride (Ber); CP-945598 HCl (CP); ezetimibe (Eze); ketoprofen (Keto); vildagliptin (Vilda).

Additionally, two compounds, CP-945598 and ezetimibe, showed trends toward increased flickering activity but did not reach statistical significance under the tested conditions, suggesting weak or condition-dependent agonistic effects (Fig.5A).

To further characterize the qualitative nature of flickering modulation, we analyzed the size distribution of flickering events. Berberine treatment not only increased the FF and RFA but also markedly shifted the size distribution of individual events across all cell lines. Specifically, berberine increased the proportion of small-size flickering events, which are considered representative of physiological mPTP interventions, while reducing the relative contribution of intermediate size events in all cell lines (Fig. 6).

**Figure 6.**
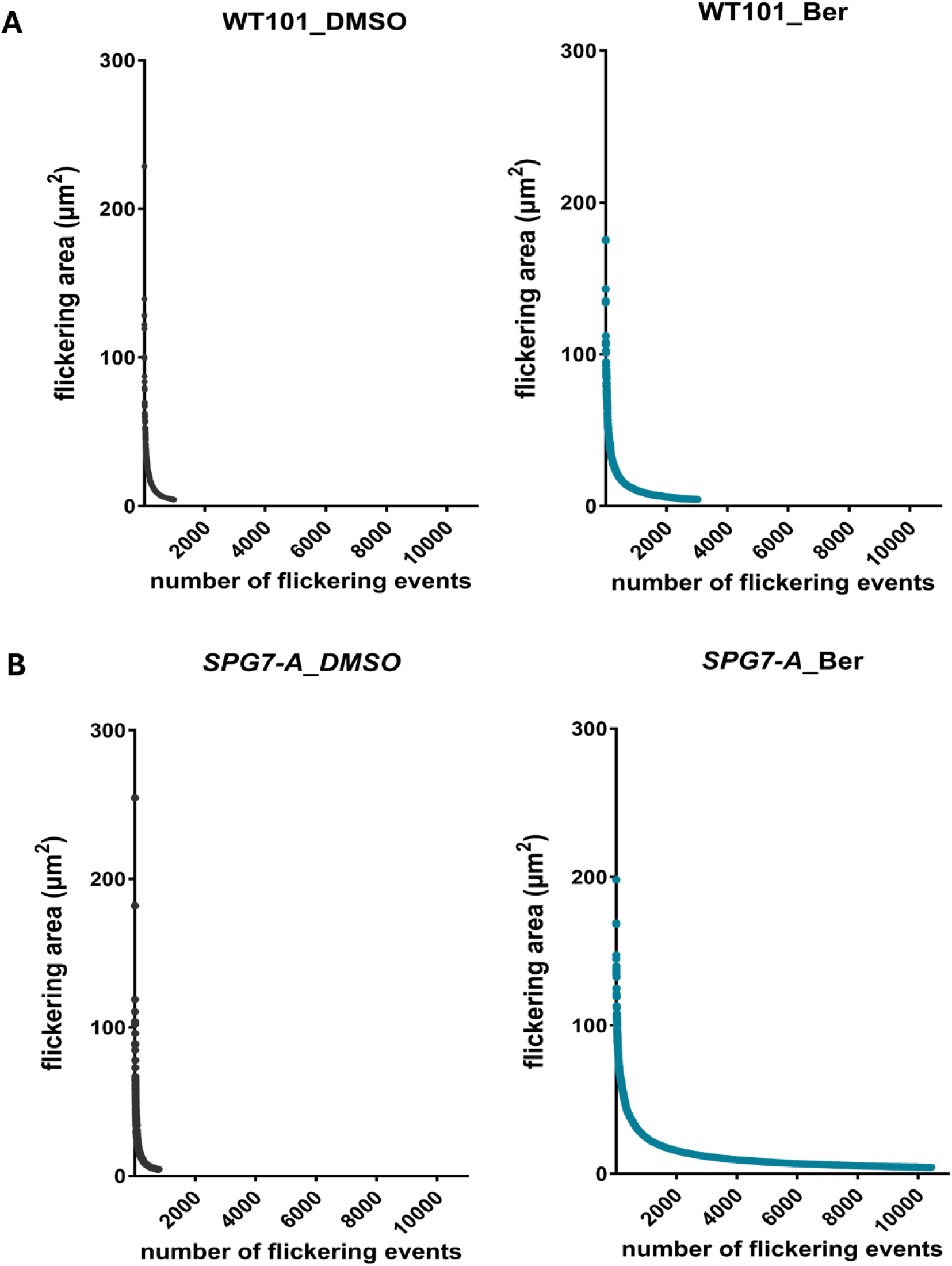
Size distribution of flickering event areas following berberine treatment. Number of events (x axis) and size distribution (y axis; values in µm^2^).

## Discussion

Uncontrolled opening of the mitochondrial permeability transition pore (mPTP) plays a pivotal role in neurodegenerative diseases, driving extensive efforts to identify effective pharmacological inhibitors [27]. Compounds such as cyclosporine A have demonstrated neuroprotective activity in preclinical models [28]. However, their clinical utility remains limited due to immunosuppressive properties and adverse effects. Additional categories, including mitochondrial antioxidants and calcium chelators, have been investigated as indirect modulators of mPTP function [29].

Despite increasing recognition of the physiological relevance of mPTP flickering, pharmacological strategies targeting the pore have remained almost exclusively focused on inhibition. To date, no clinically approved compounds exist that selectively promote controlled, low conductance mPTP opening. This represents a major therapeutic gap, particularly in conditions in which insufficient or dysregulated flickering, rather than excessive pore opening, contributes to mitochondrial dysfunction [5]. Notably, hereditary spastic paraplegia type 7 is characterized by defective mPTP flickering, which can be rescued by the agonist Bz-423 [7]. Together, these observations suggest that restoring physiological mPTP flickering through pharmacological modulation, rather than indiscriminate pore inhibition, may represent a novel and underexplored therapeutic strategy. To identify compounds capable of restoring mPTP flickering, we conducted a high-content screening of ∼2,000 FDA/EMA-approved drugs. This repurposing strategy provides advantages for rapid clinical translation while leveraging established safety profiles [9]. Using our validated TMRM/MTG-based flickering assay with automated image analysis, we systematically assessed each compound’s ability to increase both mPTP activity parameters FF and RFA.

After prioritizing candidates based on cellular and mitochondrial morphology as well as pharmacological suitability for chronic administration, 13 compounds emerged as promising modulators. In subsequent tests on fibroblasts from SPG7-A patient and a healthy control, berberine stood out as the lead candidate. Notably, berberine consistently enhanced flickering across all tested cell lines. Importantly, this effect was independent of the specific SPG7 mutation or genetic background, establishing berberine as a broadly applicable mPTP modulator rather than a mutation-specific therapeutic agent.

Berberine, an isoquinoline alkaloid derived from the *Berberis* species, demonstrated robust neuroprotective effects in numerous preclinical studies [11] [30] [31]. Recent work confirmed that berberine crosses the blood-brain barrier and enhances mitochondrial function by stimulating mitophagy and restoring mitochondrial membrane potential [32–34]. Building on these findings, our study identifies a previously unrecognized mechanism whereby berberine positively modulates physiological mPTP flickering.

Interestingly, berberine treatment not only enhanced overall FF and RFA, but also selectively boosted small- and intermediate-size flickering events. These transient, finely tuned mPTP openings are linked to mitochondrial physiological metabolic regulation through controlled release of calcium and ROS—a protective, adaptive mechanism. This stands in stark contrast to the large, sustained pore openings that drive mitochondria toward catastrophic energy dissipation. Whether berberine acts directly on mPTP structural components or indirectly through upstream mitochondrial signaling pathways remains to be determined and will require further mechanistic investigation.

Berberine has been widely employed in clinical practice to treat metabolic disorders such as type 2 diabetes, obesity, and dyslipidemia, with clinical trials consistently reporting good tolerability and a low incidence of adverse effects [35,36]

The clinical experience with berberine provides strong evidence of its safety in long-term therapeutic applications, a critical consideration for slowly progressive neurodegenerative disorders such as SPG7, as well as for other neurological and systemic diseases characterized by dysregulated mPTP dynamics, supporting its potential suitability for long-term therapeutic strategies targeting mPTP dysregulation.

## Acknowledgments

This work was supported by Fondazione Telethon ETS (grant GMR22T2019) and by the Italian Ministry of Health (RF-2019-12370417). We also gratefully acknowledge AB Medica for supporting AS. We express our gratitude to patient A for donating cells and for the continued, enthusiastic support.

---

**Supplementary Figure 1.**
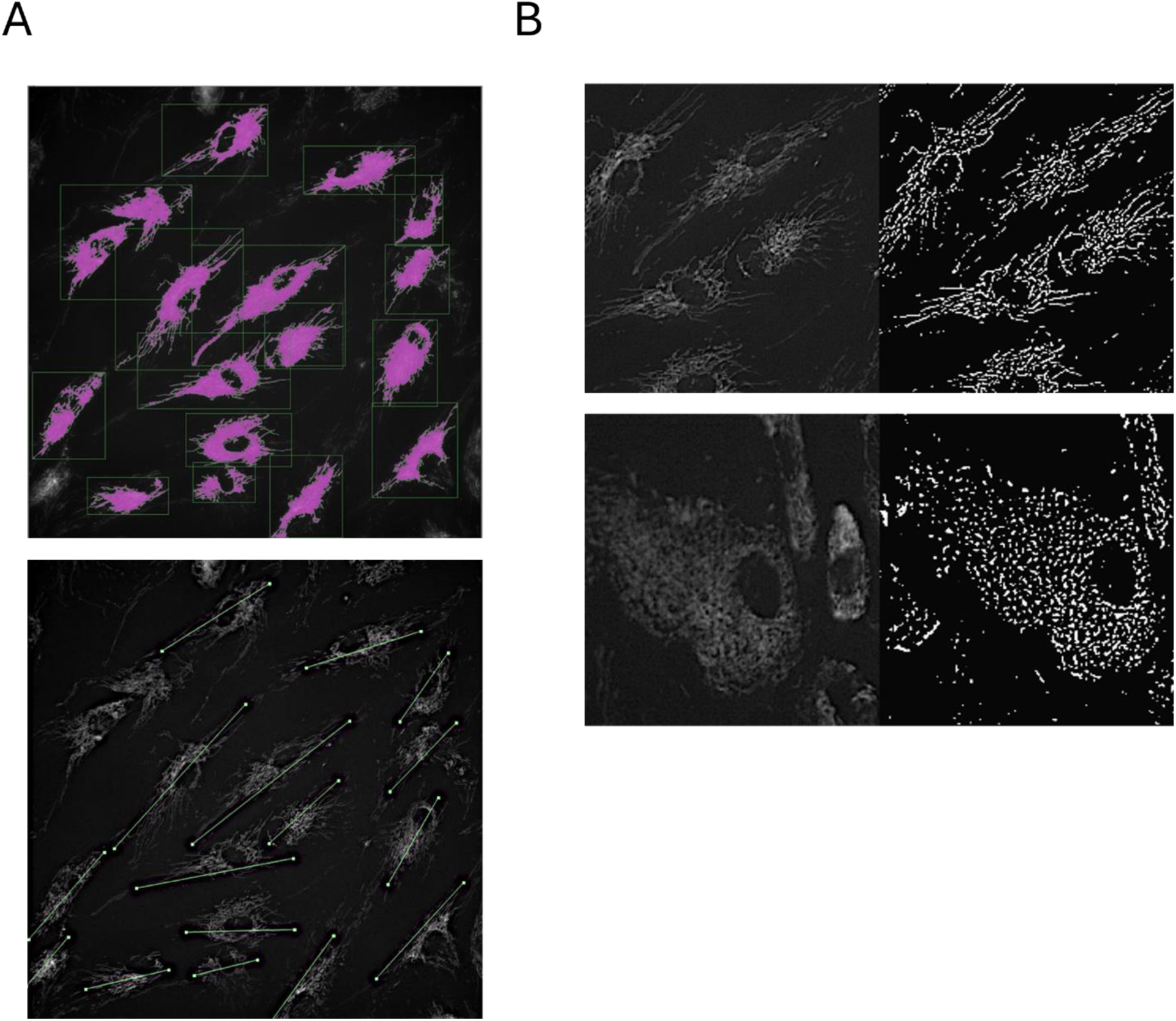
Morphological assessment of cellular and mitochondrial parameters for compound toxicity screening. (A) Volocity software pipeline output showing cellular morphology analysis. Top panel: cell shape determination for shape factor parameter. Bottom panel: longest axis measurement. Compounds were classified based on shape factor values: <0,03 indicated elongated cell morphology(healthy), >0,1 indicating rounded cell shape (cytotoxicity). Compound exhibiting shape factor >0,1 combined with reduced longest axis (value <370) were excluded as cytotoxic. (B) Mitochondrial network analysis using Mitochondria Analyzer software. Representative examples showing original fluorescence images (left panels) with corresponding mitochondrial network masks (right panels). Top panel: DMSO negative control displaying elongated mitochondria morphology; Bottom panel: test compound demonstrating altered mitochondrial network morphology compared to control. Compounds were classified based on mean form factor values: >1,6 indicating elongated, healthy mitochondria; <1,3 indicating fragmented, rounded mitochondria. Form factor values approaching 1.0 represent severely fragmented, rounded mitochondria. Compounds with form factor <1,3 were excluded due to mitochondrial toxicity

